# A novel *KCNJ16* kidney organoid model recapitulates the disease phenotype and shows restoration of lipid accumulation upon treatment with statins

**DOI:** 10.1101/2023.12.22.572750

**Authors:** E. Sendino Garví, G. van Slobbe, E.A. Zaal, J. H. F. de Baaij, J.G. Hoenderop, R. Masereeuw, M. J. Janssen, A. M. van Genderen

## Abstract

**Background:** the *KCNJ16* gene has been associated with a novel kidney tubulopathy phenotype, *viz.* disturbed acid-base homeostasis, hypokalemia and altered renal salt transport. *KCNJ16* encodes for Kir5.1, which together with Kir4.1 constitutes a potassium channel located at kidney tubular cell basolateral membranes. Preclinical studies provided mechanistical links between Kir5.1 and a disease phenotype, however, the disease pathology remains poorly understood. Here, we aimed at generating and characterizing a novel advanced *in vitro* human kidney model that recapitulates the disease phenotype to investigate further the pathophysiological mechanisms underlying the disease and potential therapeutic interventions.

**Methods:** we used CRISPR/Cas9 to generate *KCNJ16* mutant (*KCNJ16*^+/-^ and *KCNJ16*^-/-^) cell lines from healthy human induced pluripotent stem cells (iPSC) *KCNJ16* control (*KCNJ16^WT^)*. The iPSCs were differentiated following an optimized protocol into kidney organoids in an air-liquid interface.

**Results:** *KCNJ16*-depleted kidney organoids showed transcriptomic and potential functional impairment of key voltage-dependent electrolyte and water-balance transporters. We observed cysts formation, lipid droplet accumulation and fibrosis upon Kir5.1 function loss. Furthermore, a large scale, glutamine tracer flux metabolomics analysis demonstrated that *KCNJ16*^-/-^ organoids display TCA cycle and lipid metabolism impairments. Drug screening revealed that treatment with statins, particularly the combination of simvastatin and C75, prevented lipid droplet accumulation and collagen-I deposition in *KCNJ16*^-/-^ kidney organoids.

**Conclusions:** mature kidney organoids represent a relevant *in vitro* model for investigating the function of Kir5.1. We discovered novel molecular targets for this genetic tubulopathy and identified statins as a potential therapeutic strategy for *KCNJ16* defects in the kidney.

**Significance Statement:** In this study, the use of CRISPR/Cas9 technology resulted in the establishment of a *KCNJ16-*depleted kidney organoid model, instrumental in elucidating the pathophysiology of the recently reported *KCNJ16*-associated kidney tubulopathy. Our study substantiates the role of Kir5.1 (KCNJ16) in kidney disease, confirming already described phenotypes, as well as aiding to gain insight in the causal role of Kir5.1 loss in the disease phenotype. Our approach increases the knowledge on *KCNJ16*-related kidney phenotype, and it states the importance of combining CRISPR/Cas9 technology and advanced *in vitro* models for complex disease modeling and therapy testing. Furthermore, we encourage the application of our approach to the *in vitro* modeling of rare and/or underrepresented genetic kidney diseases, for which the availability of patient material is limited.

## 1 Introduction

Renal tubulopathies are a broad cluster of individual rare diseases that are mostly characterized by disruptions in cell homeostasis due to genetic defects in key renal transport proteins [1]. The *KCNJ16* gene has recently been associated with a novel kidney tubulopathy phenotype. Patients carrying biallelic loss-of-function mutations in *KCNJ16* exhibit disturbed acid-base homeostasis, severe hypokalemia, polyuria, salt craving, and Renin-Angiotensin-Aldosterone System (RAAS) activation, all of which are suggestive of salt wasting [2]. *KCNJ16* encodes for the inward rectifier potassium channel Kir5.1, which forms functional heterodimers with Kir4.2 (encoded by *KCNJ15*) and Kir4.1 (encoded by *KCNJ10*) [3]. These proteins function as potassium channels that regulate the basolateral membrane potential of several kidney epithelial cells, including those lining the PT (PT), cortical thick ascending limb (cTAL), distal convoluted tubule (DCT), connecting tubule (CNT), and collecting duct (CCD) [4, 5]. Altered membrane potential through dysfunctions of Kir4.1/Kir5.1 or Kir4.2/Kir5.1 heteromeric channels leads to dysregulation of several membrane potential-dependent transport processes and protein activities [6, 7], resulting in a tubulopathy phenotype as observed in patients with *KCNJ16* loss-of-function [2, 3]. In the PT, Kir4.2/Kir5.1 channels regulate the membrane potential in response to changes in intracellular pH. When the intracellular pH decreases, Kir4.2/Kir5.1 channels are activated and depolarize the cell membrane, thereby inhibiting the Na-bicarbonate cotransporter 1 (NBC1; *SLC4A4*) that facilitates bicarbonate reabsorption from the PT cells into the circulation. This, ultimately, results in a hyperpolarization of the membrane and activation of NBC1 [8–10]. Dysfunction of Kir4.2/Kir5.1 leads to metabolic acidosis, accompanied by an increase in intracellular pH, membrane depolarization and reduced NBC1 activity in the PT [9, 11]. These findings suggest that the feedback loop between pH, Kir4.2/Kir5.1 and NBC1 activity guards acid-base homeostasis. In the DCT, Kir5.1 acts both as a potassium transporter as well as an extracellular potassium sensor involved in the potassium conductance of the basolateral membrane of the cells. Upon sensing of the extracellular potassium concentration, the channel changes the membrane potential and adjusts the intracellular chloride concentration [12, 13]. Furthermore, Kir4.1/Kir5.1 channels showed to be involved in sodium reabsorption and potassium secretion in the CD, where sodium reabsorption is facilitated by the epithelial Na^+^ channel (ENaC) [14] and potassium secretion is facilitated by the outer medullary potassium channel (ROMK) [15], which is considered one of the main routes of potassium secretion in the kidney.

Overall, previous evidence suggests that defects in potassium transporters such as *KCNJ10* and *KCNJ15* is associated with a kidney phenotype. However, there is no precedent for *KCNJ16* defects alone causing such a phenotype except for the recently published work by Schlingmann *et al.* [2], whereas other studies describing the involvement of *KCNJ10* and *KCNJ15* in kidney disease were mostly based on rodent models. This subscribes the need for translational models to better understand the mechanism of disease in humans and to study therapeutic interventions. Human kidney organoids are considered advanced *in vitro* models that could potentially bridge the knowledge gap. Recent advances in this field demonstrated that human induced pluripotent stem cell (iPSC)-derived kidney organoids form highly organized, polarized, 3D structures containing most cell types of the nephron, and respond to environmental cues [16–19]. Moreover, iPSC-derived kidney organoids combined with CRISPR/Cas9 technology have shown to be a powerful tool to study genetic kidney disorders by recapitulating complex pathological markers/processes such as cyst formation in polycystic disease [20–22]. Therefore, the generation of an advanced, human-derived *in vitro* model that recapitulates aspects of the phenotype offers a tool to further study the disease and to create a personalized, pharmacotherapeutic platform for intervention studies.

In this study, we generated *KCNJ16* knock-out human iPSC-derived kidney organoids to provide an advanced and more human-translatable model to study the underlying pathophysiological mechanisms of Kir5.1 dysfunction. Additionally, we used this system to further unravel potential disease-causing mechanisms and to evaluate the effect of several drug compounds as novel therapeutic options for kidney disease caused by defects in *KCNJ16*.

## 2 Results

### 2.1 Gene editing using CRISPR/Cas9 in kidney organoids results in loss of Kir5.1

To assess whether iPSC-derived kidney organoids were a suitable platform to model and study *KCNJ16* loss, the mRNA and Kir5.1 protein were measured (Figure 1). Our results show an increase in *KCNJ16* mRNA expression during iPSC differentiation and maturation into kidney organoids (Figure 1B), suggesting that *KCNJ16* expression is relevant for kidney differentiation. Moreover, a comparison of *KCNJ16* mRNA expression between mature organoids and a healthy kidney biopsy (Figure 1B) revealed similar levels, indicating that mature kidney organoids represent a relevant *in vitro* model for investigating the function of Kir5.1.

**Figure 1.**
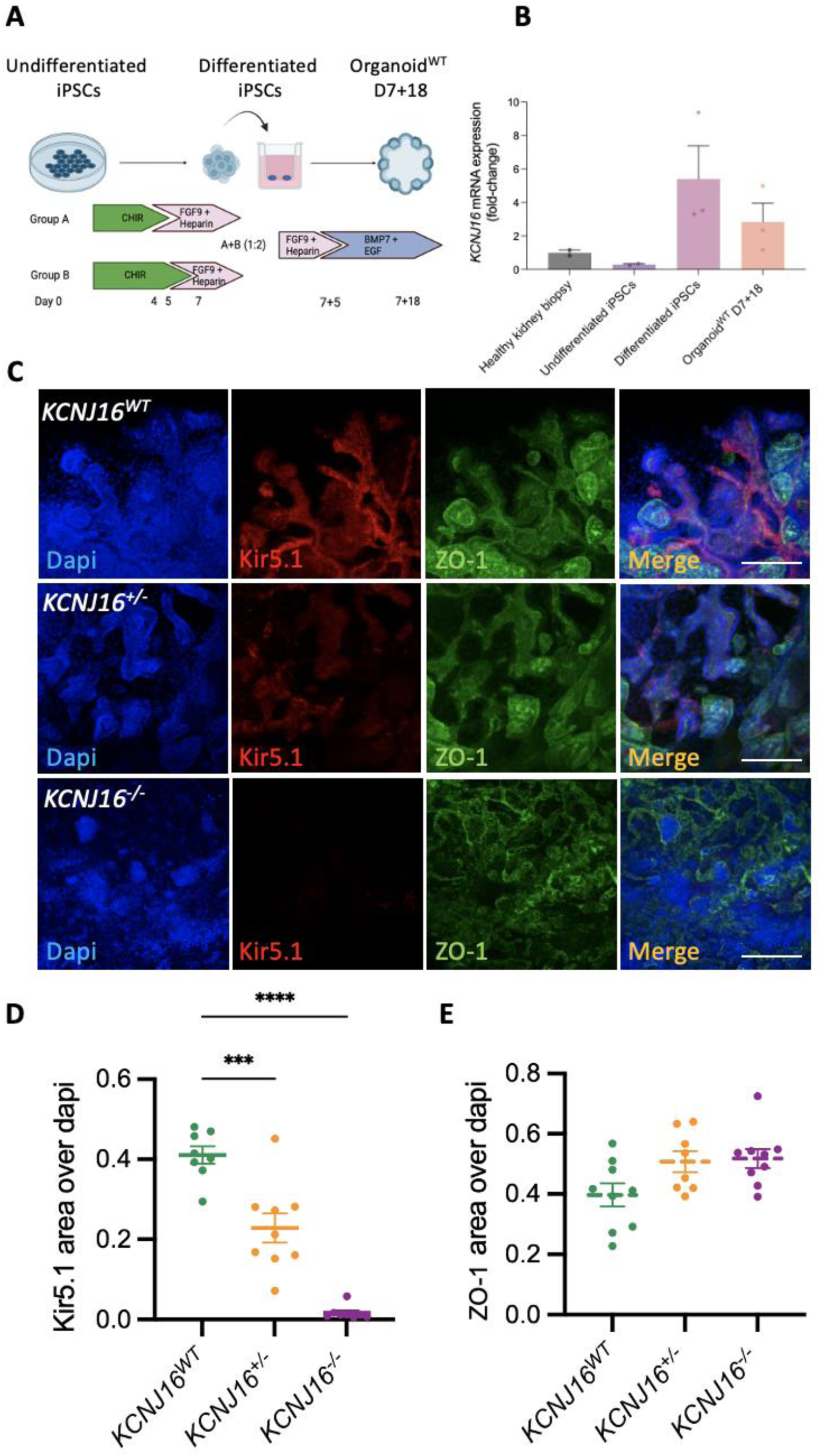
Human iPSC-derived *KCNJ16*-depleted kidney organoids show Kir5.1 depletion. (**A)** Schematic presentation of the differentiation protocol from undifferentiated iPSCs to mature kidney organoids. (**B)** The mRNA expression levels of *KCNJ16* in undifferentiated iPSCs and mature organoids compared to the levels of a healthy human kidney biopsy sample. (**C)** Representative immunofluorescence images of Kir5.1 in *KCNJ16^WT^*, *KCNJ16^+/-^* and *KCNJ16^-/-^*organoids, in conjunction with the tight-junction marker zonula occludens-1 (ZO-1). (**D)** Semi-quantification of the partial and total loss of Kir5.1 (*KCNJ16*) in *KCNJ16^+/-^* and *KCNJ16^-/-^* compared to *KCNJ16^WT^*, respectively. (**E)** Semi-quantification of ZO-1 staining with similar expressions in all three *KCNJ16* organoid lines. Scale bar represents 100µm. Statistics represent the significance of one-way ANOVA test (n=3, ****= p<0.0001, ***= p<0.001).

Next, we used CRISPR/Cas9 to generate *KCNJ16* knock-out iPSC lines. After single cell sorting, iPSC colonies were genotyped and both a clone with a homoallelic mutation (*KCNJ16^+/-^*) and a clone with a biallelic mutation (*KCNJ16^-/-^*) in the exon 5 of the *KCNJ16* gene were selected for further analysis. Immunostaining showed that Kir5.1 (*KCNJ16*) was expressed at the basolateral membrane of tubular cells, with tight junctions (ZO-1) located between cells at the apical side in control organoids containing wild type Kir5.1 (*KCNJ16^W^*^T^) (Figure 1C). On the contrary, both *KCNJ16^+/-^*and *KCNJ16^-/-^* organoids showed a partial (46%) and complete (99%) depletion of Kir5.1 expression, respectively. Despite this, the abundance of ZO-1 protein seemed unaffected by the loss of Kir5.1 but served as a guide to delineate the kidney organoids tubular structures (Figure 1C-E).

### 2.2 Loss of Kir5.1 results in acid/base homeostasis imbalance and altered ion transport

To investigate the impact of Kir5.1 loss in kidney organoids, we evaluated the transcriptional landscape of *KCNJ16^-/-^* compared to *KCNJ16^WT^* which revealed 3,519 differentially expressed genes (1,963 up and 1,556 down) (Figure 2A-B). We clustered the significant genes by their molecular function which shows that upon loss of Kir5.1, voltage- and charge- dependent ion transporters were upregulated while DNA- and RNA-binding genes were found downregulated (Figure 2C). Complementary, we investigated the network representation of the GAGE pathway analysis after clustering by molecular function, which confirmed that loss of Kir5.1 led to the upregulation of various cell membrane ion/cation channels and transporters dependent on charge and/or cell membrane voltage (Figure 2D). Further exploration using KEGG pathway analysis confirmed alterations in several key transcripts involved in the cAMP pathway (Figure S1), PT bicarbonate reclamation (Figure S2), and mineral reabsorption (Figure S3) upon Kir5.1 loss.

**Figure 2.**
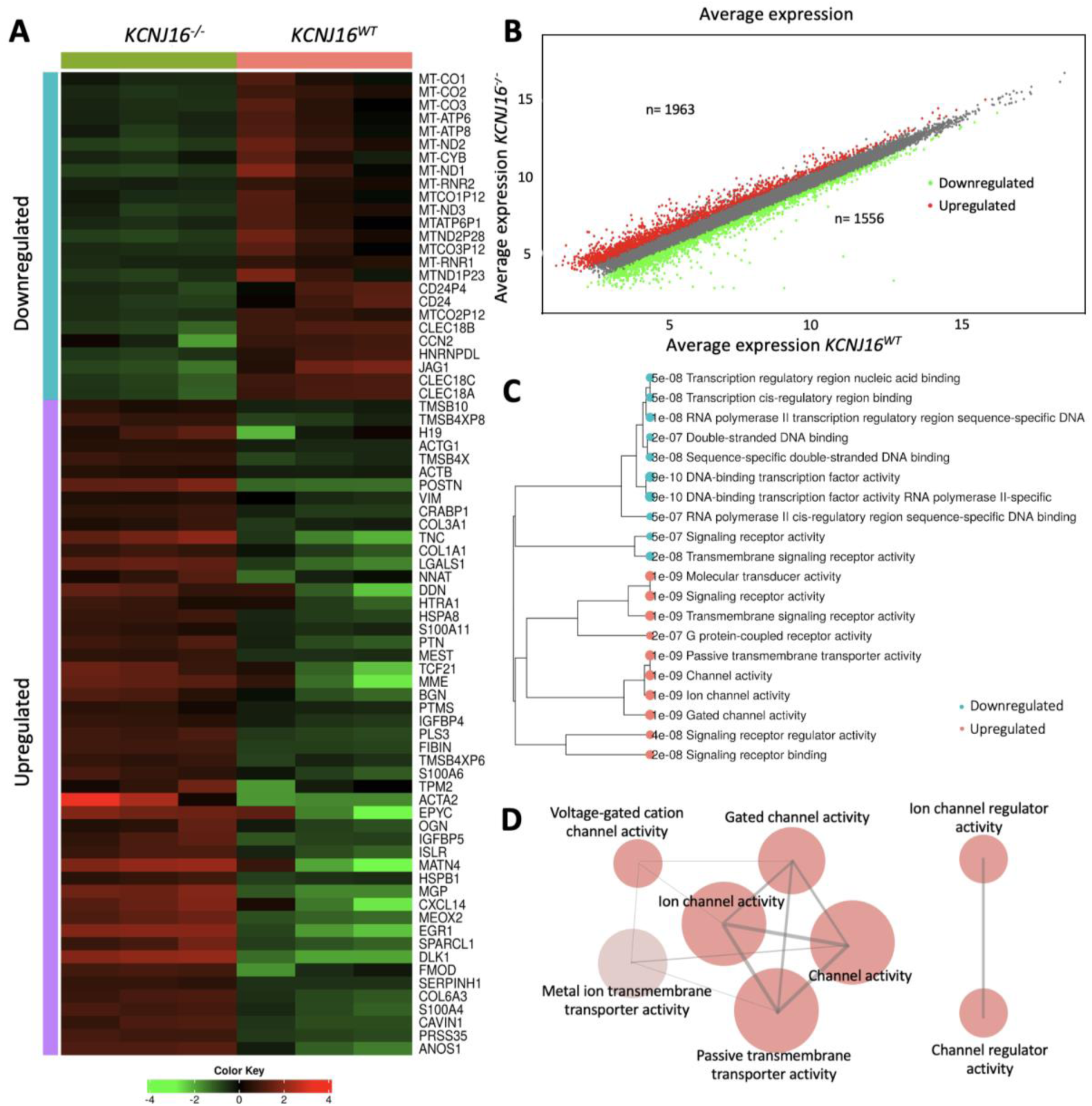
RNA-Seq analysis reveals alterations in ion- and electrolyte transporters upon loss of Kir5.1. (**A)** Heatmap representation of the hierarchical clustering (Pearson, p<0.0001) including the top 100 most differentially expressed genes between *KCNJ16^WT^* and *KCNJ16^-/-^*. (**B)** Scatter plot presenting the total number of up- and down- regulated genes when comparing *KCNJ16^-/-^* to *KCNJ16^WT^* (fold change cut off: 1.5, p<0.0001). (**C)** Clustering tree presentation of the DEG2 analysis after sorting the most differentially expressed genes by molecular function in *KCNJ16^-/-^*. (**D)** Network presentation of the GAGE pathway analysis showing the most enriched pathways in *KCNJ16^-/-^* sorting by molecular function (FDR cutoff: 0.5). In the network plots (**D**), the size of the nodes represents the number of genes enriching that pathway (gene set minimum size:10). The red color represents upregulated pathways, and the opacity of the nodes represents the fold enrichment (p<0.001). Two pathways (nodes) are connected if they share at least 30% of genes.

To investigate the effect of the potential ion- and electrolyte transport impairments, as well as the cAMP pathway alterations hinted by the transcriptomic data, we first evaluated the ability of the kidney organoids to respond to forskolin treatment (Figure 3). In untreated conditions, all nephron segments could be qualitatively observed in all three lines, but we detected the presence of enlarged tubular structures (particularly proximal and distal tubules) in the *KCNJ16*^+/-^ and *KCNJ16*^-/-^ that were absent in *KCNJ16*^WT^ kidney organoids (Figure 3A-B). Upon treatment with forskolin, we observed the formation of cysts (mainly in the proximal and distal tubular structures) in both *KCNJ16*^+/-^ and *KCNJ16*^-/-^ (Figure 3C-D), which were higher in number when compared to the *KCNJ16^WT^* kidney organoids (Figure 3E).

**Figure 3.**
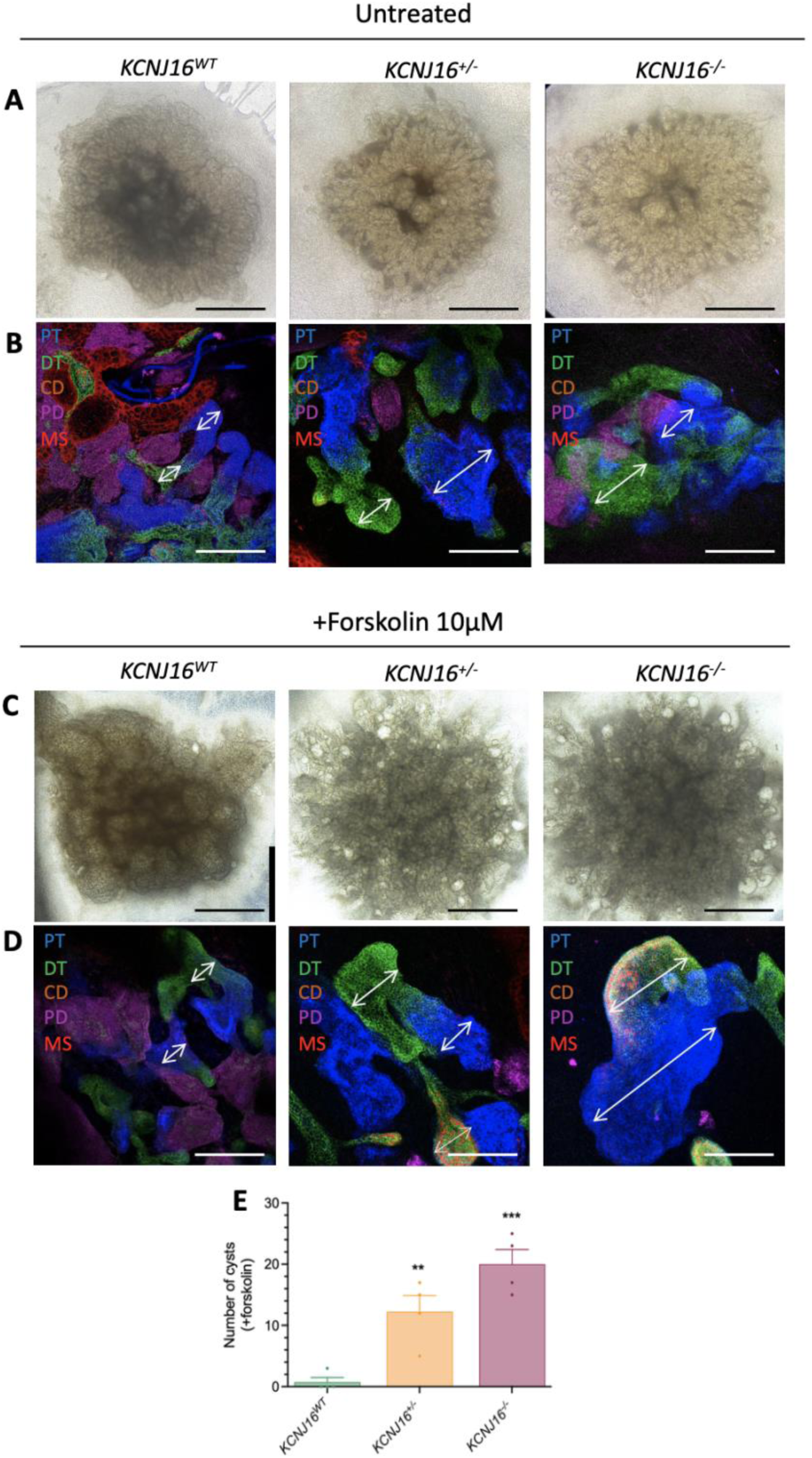
Partial and total loss of Kir5.1 results in cyst formation upon forskolin treatment. **(A)** Brightfield images of all three *KCNJ16* organoids without forskolin treatment. (**B)** Immunofluorescent images at of all three *KCNJ16* organoids without forskolin treatment. (**C)** Representative brightfield images of all three *KCNJ16* organoids after 24h treatment with 10µM forskolin. (**D)** Immunofluorescent images of all three *KCNJ16* organoid lines upon 24h treatment with 10µM forskolin. (**E)** Semi-quantification of the cyst formation. PT: proximal tubule, DT: distal tubule, CD: collecting duct, PD: podocytes, MS: mesangial cells. Scale bar represents 100µm. Statistics represent the significance of one-way ANOVA test (n=3, ***= p<0.001, **= p<0.01).

The cyst formation upon forskolin treatment indicates either a malfunction of the cAMP-PKA pathway itself, as supported by our RNA-seq data (Figure S4A), and alterations in downstream players responsible for water transport. To further investigate this, *KCNJ16*^-/-^ and *KCNJ16*^WT^ kidney organoids were treated with a combination of forskolin and an inhibitor to block either aquaporin-1 (tetraethylammonium; TEA), inhibit urea transport (acetamide), or to block Na^+^-K^+^-ATPase (digoxin) (Figure S4B-C). We observed an increase in cyst formation in the *KCNJ16*^WT^ treated with a combination of forskolin and acetamide when compared to forskolin treatment alone, while the same treatment reduced cyst formation in *KCNJ16*^-/-^ (Figure S4C). Furthermore, treatment of *KCNJ16*^WT^ with the combinations of forskolin and digoxin or forskolin and TEA hinted towards a mild inhibition of cyst formation when compared to forskolin treatment alone, comparable with *KCNJ16*^-/-^ for the same comparisons (Figure S4C). Overall, these results indicate an increased sensitivity to water imbalances upon Kir5.1 loss, especially with forskolin as stressor. Additionally, the increase in cysts formation in *KCNJ16*^WT^ together with the improvement in cysts formation in *KCNJ16*^-/-^ after adding acetamide might suggest the role of urea transport as potentially contributing to the water transport alterations.

Secondly, we evaluated the ability of the kidney organoids to regulate the intracellular pH upon stress (Figure 4). Under normal conditions, the intracellular pH of all three organoids lines was similar (Figure 4A). Upon NaHCO_3_ addition, the intracellular pH of all organoid lines increased and the response of both *KCNJ16*^+/-^ and *KCNJ16*^-/-^ was comparable to that observed in *KCNJ16*^WT^. However, *KCNJ16*^-/-^ failed to restore intracellular pH after 12 minutes, while within that time frame *KCNJ16*^WT^ organoids partially restored their intracellular pH upon 5 minutes of extracellular NaHCO_3_ stress. Additionally, *KCNJ16*^+/-^ organoids showed a large variability in their response to pH stress, consistent with a more intermediate phenotype. To further understand the effect of Kir5.1 loss on acid/base homeostasis, we quantified the mRNA of key transporters involved in bicarbonate handling (Figure 4B). Both *KCNJ16*^WT^ and *KCNJ16*^-/-^ were able to increase their intracellular pH upon NaHCO_3_ addition, indicating that the transporters that allow the entry of NaHCO_3_ (such as *NBC1* and *NBCe2*) were not affected by Kir5.1 loss, while the transporters in charge of responding to a NaHCO_3_-induced acid/base stress (such as *CA4*, *SNAT1* and *SNAT3*, *NHE3*, and *PCK1*) showed altered expression levels (Figure 4B). Furthermore, our RNA-Seq data showed an overall downregulation of key genes (including WNK1/2/4, ATP1A1, FXYD2, and ENaC) that could also play a role in acid/base homeostasis via their compensatory mechanisms upon NaHCO3 stress in the distal tubule and collecting duct (Figure S5).

**Figure 4.**
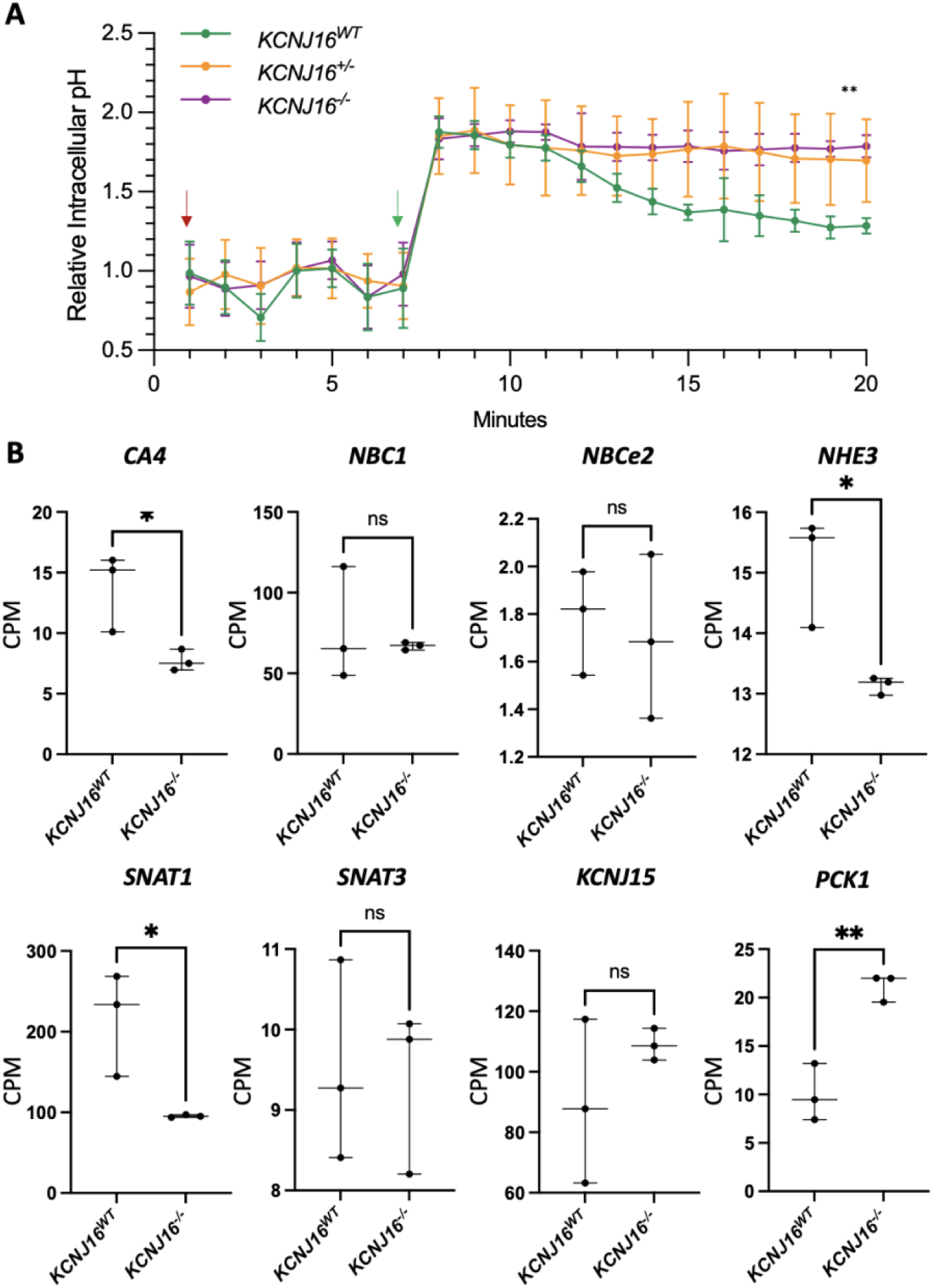
Loss of Kir5.1 results in the inability to regulate intracellular pH upon NaHCO_3_ stress. **(A)** Relative intracellular pH values before and after the addition of NaHCO_3_ in the culture media. The red and green arrows indicate the time of the first measurement after adding the pH probe and after adding 10 mM NaHCO_3_, respectively. (**B)** The mRNA levels quantified as count per million (CPM) of key genes involved in bicarbonate handling in the kidney. Statistics in panel A represent the significance of one-way ANOVA test, while statistics in panel B represent unpaired t-test (N=3, ***= p<0.001, **= p<0.01).

#### *KCNJ16* depleted kidney organoids show altered TCA cycle and lipid accumulation

To further evaluate the downstream effect of the acid/base homeostasis disruption upon Kir5.1 loss and the ion- and electrolyte transport impairment, a metabolomic analysis was performed. While the metabolic landscape of *KCNJ16*^+/-^ and *KCNJ16*^WT^ partly overlapped, the metabolic pattern of *KCNJ16*^-/-^ clustered individually and distinct (Figure 5A). The heatmap of differentially expressed metabolites revealed that the differences between *KCNJ16*^WT^ and *KCNJ16*^-/-^ were mostly due to differential accumulation of short chain acyl-carnitines, aminoacids, and intermediates of the TCA cycle and glycolysis (Figure 5B). Subsequently, we found that the metabolic pathways driving the major differences between *KCNJ16*^WT^ and *KCNJ16*^-/-^ were related to amino acids, TCA cycle, and glycolipid metabolism (Figure 5C).

**Figure 5.**
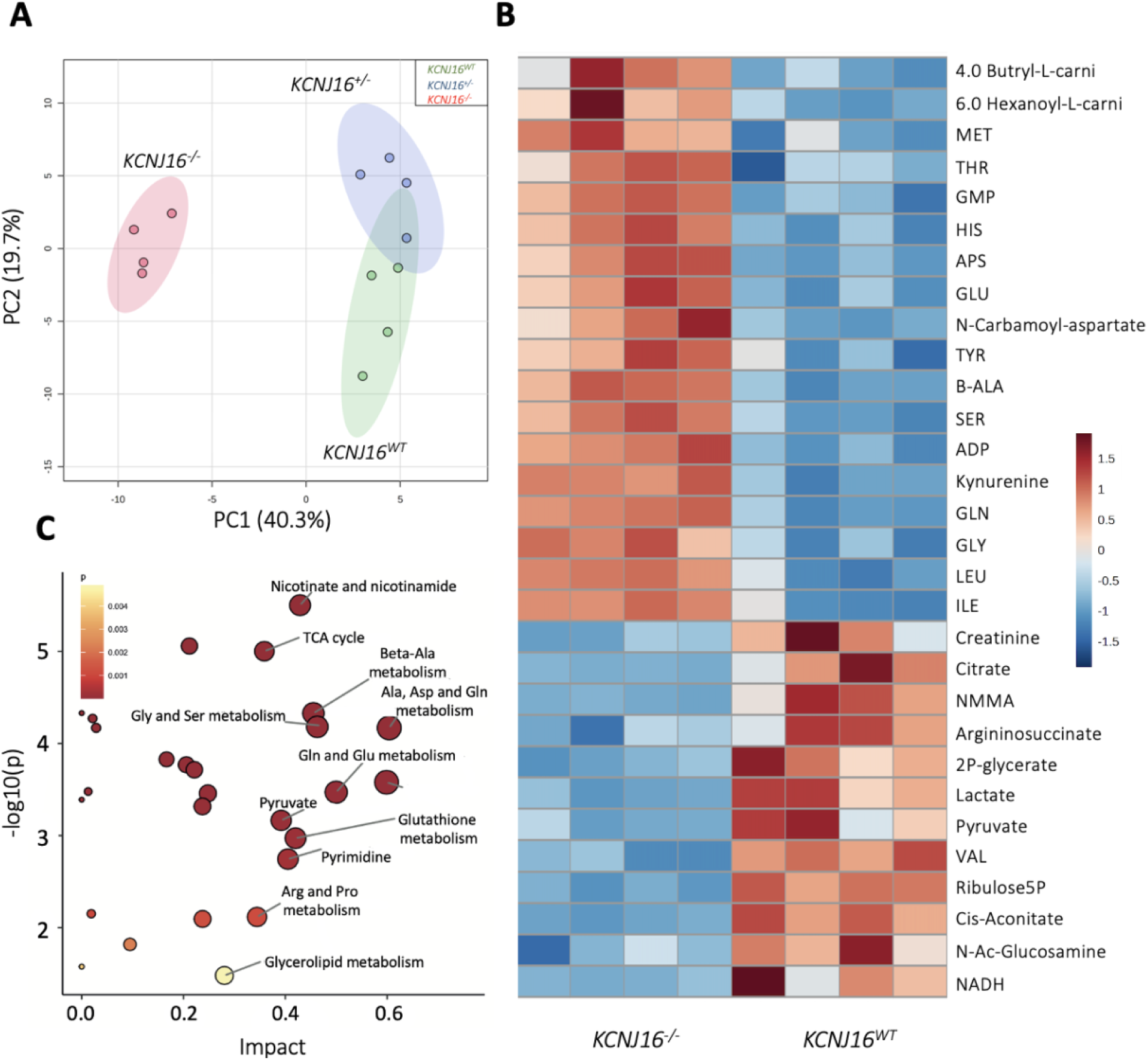
Metabolomics analysis shows the impact of the total loss of Kir5.1 on key metabolic pathways. (**A)** Principal component analysis (PCA) plot showing the differences between *KCNJ16*^-/-^ and both *KCNJ16*^WT^ and *KCNJ16*^+/-^, as well as the similarities between *KCNJ16*^WT^ and *KCNJ16*^+/-^, in their metabolic landscape. (**B)** Heatmap of the 30 most differentially expressed metabolites when comparing *KCNJ16*^WT^ and *KCNJ16*^-/-^. (**C)** Graphical presentation of the most differentially enriched pathways between *KCNJ16*^WT^ and *KCNJ16*^-/-^.

To further explore the suggested TCA cycle dysfunction, we performed a metabolic flux study using ^13^C-labeled glutamine to follow the incorporation of glutamine-derived carbons in TCA cycle intermediates (Figure 6). In terms of carbon-labelled metabolites, the relative ^13^C-labeling indicated the accumulation of glutamine and glutamate in the *KCNJ16*^-/-^ kidney organoids but no relevant differences were found in the glutamine usage within the TCA cycle. Total quantification of the TCA cycle intermediates showed a slight accumulation of α-ketoglutarate (α-KG) and a generalized downregulation of the other metabolites, especially cis-aconitate and citrate in *KCNJ16*^-/-^ kidney organoids, indicating low TCA cycle activity. Additionally, *KCNJ16*^-/-^ organoids showed a reduction in pyruvate and acetyl-CoA, which together with the low levels of citrate might be indicative of the usage of the TCA intermediates towards fatty acid synthesis.

**Figure 6.**
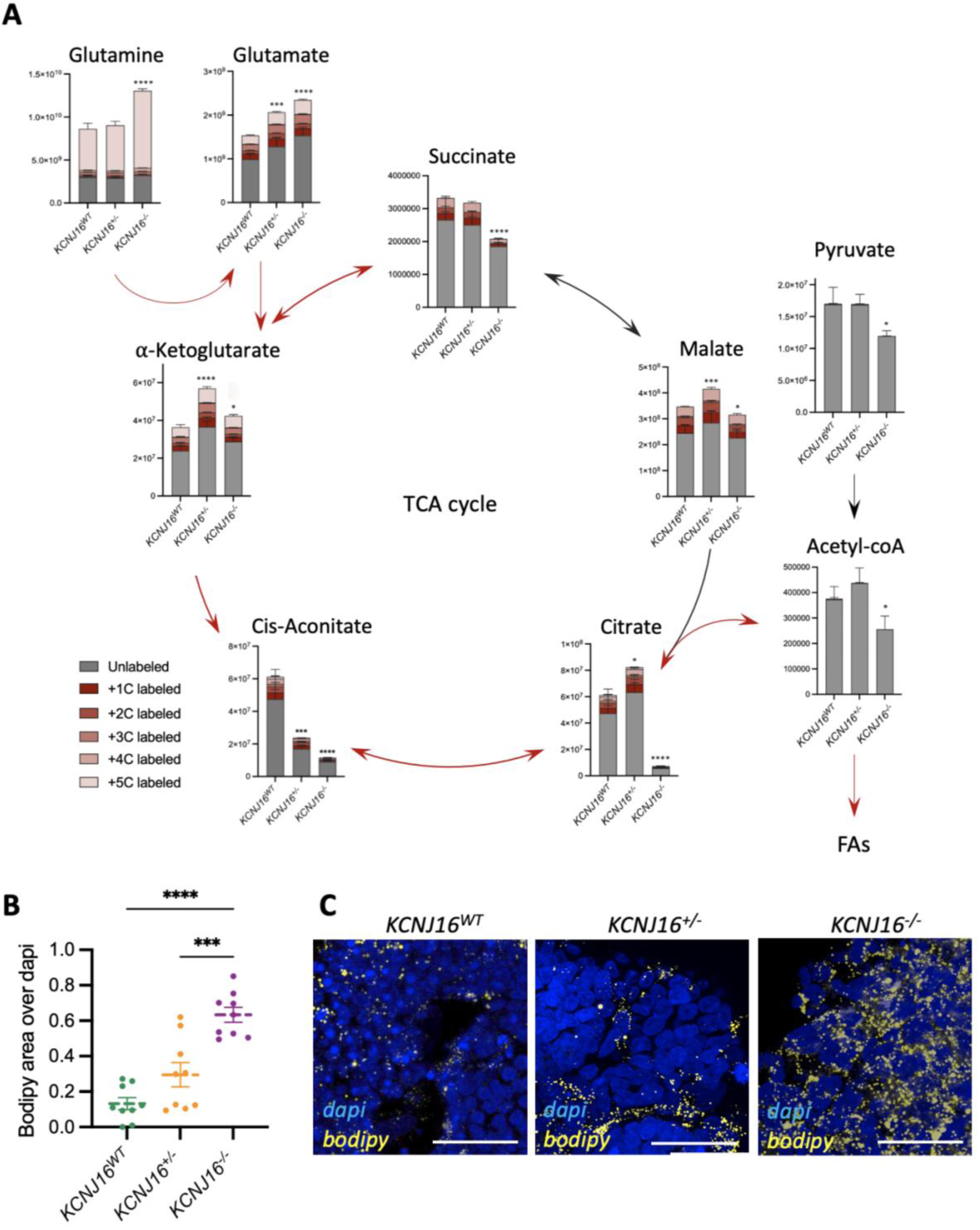
Loss of Kir5.1 results in TCA cycle impairments and lipid droplet accumulation. (**A)** Individual quantification of the main metabolites of the TCA cycle, including different isotopes with ^13^C. Red arrows represent the route within the TCA cycle that results in fatty acid synthesis. (**B**) Semi-quantification of lipid droplet accumulation using bodipy as a probe. (**C**) Representative immunostainings showing the lipid droplet accumulation using bodipy as a marker. Scale bar 10µm. Statistics represent the significance of one-way ANOVA test (n=4, ****= p<0.0001, ***= p<0.001, *= p<0.05) of the total metabolites amount compared to the *KCNJ16*^WT^ control values.

To evaluate this, we analyzed the presence of lipid droplets in all three kidney organoid lines (Figure 6). *KCNJ16*^-/-^ organoids clearly demonstrate accumulation of lipid droplets as compared to *KCNJ16*^WT^, while the heterozygous organoids show an intermediate phenotype (Figure 6 B-C). The accumulation of lipids was accompanied by an increased deposition of extracellular matrix proteins, as evaluated by collagen-I and fibronectin (Figure S6). In kidney failure, deposition of matrix proteins is observed during fibrosis for which TGFß is considered a central mediator. Indeed, exposure to the growth factor led to increased collagen-I accumulation in *KCNJ16*^WT^ and *KCNJ16*^+/-^ organoids, whereas *KCNJ16*^-/-^ were not responsive to TGFß stimulation (Figure S6). Furthermore, after treatment with TGFß, fibronectin levels increased but only in *KCNJ16*^WT^, and to the level observed in *KCNJ16*^-/-^ and *KCNJ16*^+/-^ with and without stimulation.

#### Statins prevent lipid accumulation and fibrosis phenotype in *KCNJ16*-depleted kidney organoids

Finally, we investigated whether the lipid droplet accumulation and matrix proteins deposition could be reversed by treating *KCNJ16*^-/-^ organoids with the statins simvastatin or pravastatin, and a fatty acid synthase blocker (C75), alone and in combinations (Figures 7 and S7). Since we observed that lipid droplets started to accumulate in *KCNJ16*^-/-^ within the last 6 days of maturation, organoids were exposed daily to the drugs starting from d7+12. Statins and C75 all effectively reduced lipid droplet accumulation in *KCNJ16*^-/-^ organoids to matured, untreated *KCNJ16*^WT^ organoid levels (Figure 7 A-B). Furthermore, statins and C75 reduced collagen I deposition alone and in combination, with the combination being most effective (Figure S7). In contrast, this effect was not observed for fibronectin deposition. These findings suggest that statins and a fatty acid synthase blocker partially reverse the phenotype associated with Kir5.1 deficiency.

**Figure 7.**
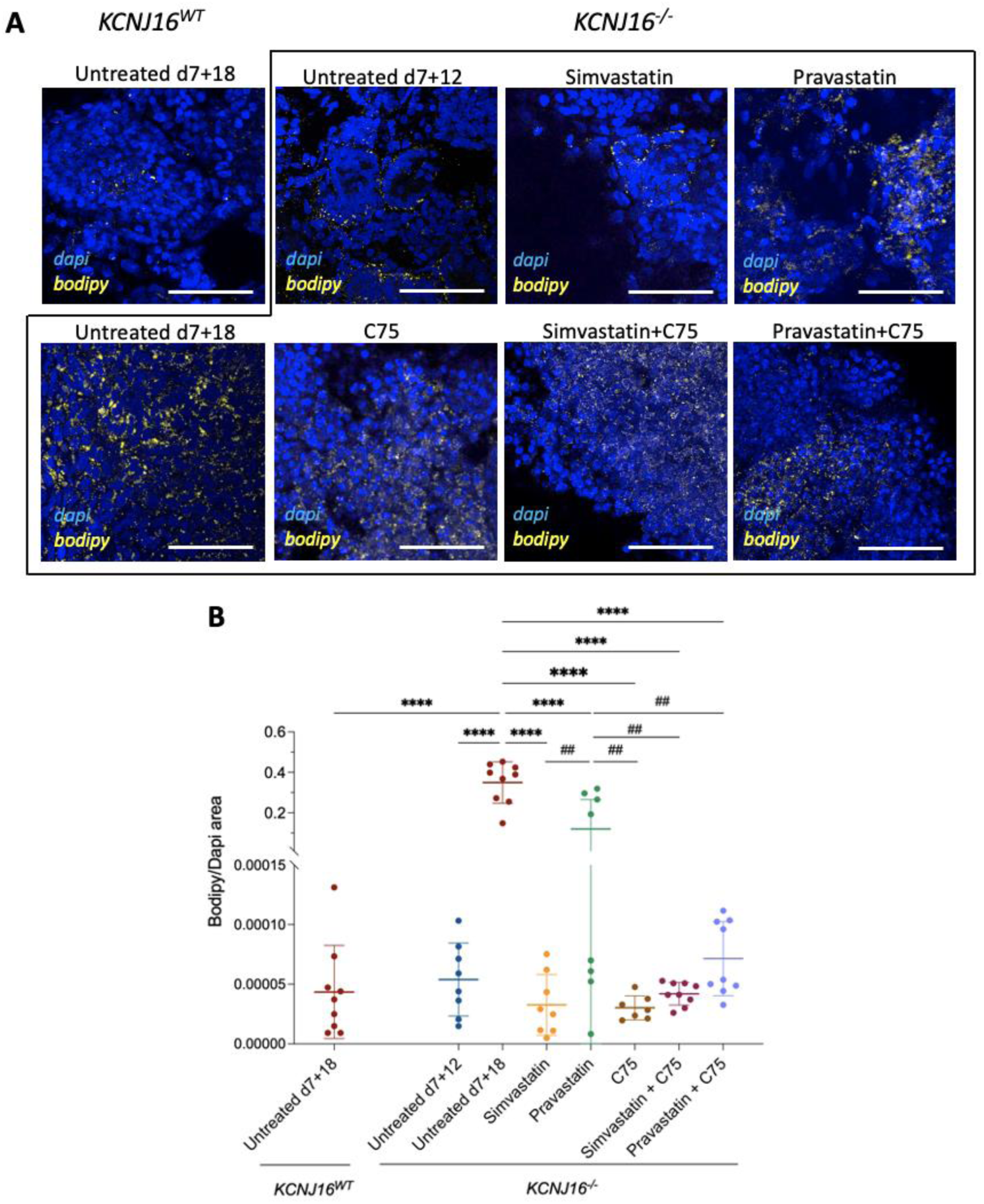
Treatment with statins prevents lipid droplet accumulation in *KCNJ16*^-/-^ kidney organoids. (**A)** Representative immunofluorescence images of lipid droplets in *KCNJ16*^-/-^ organoids after 7 days treatment with Simvastatin, pravastatin, C75, and combinations. (**B)** Semi-quantification of droplet accumulation imaging data. Simvastatin and pravastatin were used at a final concentration of 0.5 µM and C75 was used at a concentration of 40 µM. Scale bar 100µm. Statistics represent the significance of one-way ANOVA tests (n=3, ****= p<0.0001 when comparing all conditions to the untreated *KCNJ16*^-/-^ at d7+18, and ^##^= p<0.01 when comparing treatments amongst each other).

## Discussion

In this study, we aimed to generate and characterize a novel *KCNJ16* knock-out kidney organoid model to provide an advanced and robust platform to further study this recently described kidney tubulopathy [2]. Our results show that total loss of of Kir5.1 (*KCNJ16*) led to transcriptomic impairment of key transporters regulating essential cellular homeostasis and metabolic processes, which resulted in pH imbalance, cyst formation and metabolic landscape impairment, with particular impact on the TCA cycle and lipid metabolism. Subsequently, the depletion of Kir5.1 resulted in lipid droplet accumulation and fibrosis, which was successfully prevented by treatment with statins.

Along the different segments of the nephron, Kir5.1 can form dimers with Kir4.1 and Kir4.2. While the last two can also form dimers with themselves, Kir5.1 cannot [23]. Previous studies showed that the loss of Kir4.1 and/or Kir4.2 in Kir5.1 dimer lead to the impairment of channel function, resulting in structural damage and impaired salt reabsorption in the distal tubule, hypocalciuria, hypomagnesemia, and hypokalemic acidosis in the kidney of mice [24–28]. Supporting evidence of the effects of only Kir5.1 depletion remains limited because of a lack of robust *in vitro* and *in vivo* models and limited clinical cases. Our results show that upon Kir5.1 depletion, the activity of ion, electrolyte, salt and water transporters is impaired, reducing the ability of *KCNJ16*^-/-^ organoids to regulate their water transport and intracellular pH upon external stimuli. This is also in line with transporter alterations upon Kir5.1 loss observed at transcriptomic level (Figure 4, S4 and S5). Membrane potential and pH imbalances have been directly linked to electrolyte homeostasis and water balance impairment, especially in the kidney [29–31], which could potentially lead to water accumulation and cyst formation, as observed in our *KCNJ16*^-/-^ kidney organoids. Although the specific role of Kir5.1 in water/electrolyte balance and pH regulation is yet unknown, our results showed that loss of function of KCNJ16 may alter the same feedback loop between water/electrolyte transport and pH homeostasis which could eventually result in the phenotype observed in patients, including salt wasting, hypokalemia and metabolic acisosis [2]. The cyst seen in our organoid model are indicative of abnormal water transport into the lumen of closed tubular structures and do not suggest patients may develop kidney cysts. We therefore used the cyst formation as a readout for water/electrolyte transport. To further understand the inability to regulate water transport upon forskolin stress, we pharmacologically inhibited the transport of sodium, potassium, water, bicarbonate and urea. Our results indicate that urea transport might be alteres upon Kir5.1 loss resulting in cyst formation. However, these results must be taken with caution because of a lack of specificity of the inhibitors used [32–35].

Another key feature identified upon Kir5.1 loss is the impairment in the TCA cycle and lipid metabolism. The defects we detect in the TCA cycle could be due to changes in the activity of enzymes that regulate either the TCA cycle itself or that are involved in the synthesis and processing of the TCA cycle precursors, such as citrate synthase, aconitase and pyruvate dehydrogenase. The impaired functioning of these enzymes could directly lead to disruptions in glucose and glutamine metabolism, which could in turn affect the functioning of the TCA cycle by limiting the presence of key sources. Our transcriptomic data did not show significant differences in these enzymes; however, their activity might still be impacted, since these enzymes are directly affected by alterations in the acid/base environment they are in. Furthermore, potential changes in activity of enzymes that regulate the amino acids and lipid metabolism could either contribute further to the observed impairments in metabolic pathways or be a consequence of it, given that these pathways change directionality depending on the energy demands of the cells. Citrate showed to be the most differentially expressed metabolite within the TCA cycle when comparing *KCNJ16*^WT^ and *KCNJ16*^-/-^ organoids. This metabolite is involved in several metabolic pathways [36, 37] and its observed loss in *KCNJ16*^-/-^ could have different implications in pathways in addition to the TCA cycle [38]. Citrate can be either processed further within the TCA cycle directly into succinate, but it can also enter a different loop outside the TCA cycle towards lipid synthesis, depending on the energy demands and the anabolic/catabolic status of the cells [39–41]. Our results lead us to hypothesize that citrate exits the TCA cycle to enter the lipid biosynthesis pathway, a phenomenon that has been observed in multiple diseases, including cancer [42] and diabetic nephropathy [43]. Furthermore, it has been shown previously that low conversion of phosphoenolpyruvate (PEP) into pyruvate can lead to the accumulation of glycolytic intermediates [44], which can then be used for either nucleotide synthesis and/or lipid synthesis [45]. Although we observe reduced levels of pyruvate in *KCNJ16*^-/-^, our results showed lower levels of glycolytic intermediates, as well as a lower presence of TCA intermediates leading towards fatty acid synthesis, such as citrate and acetyl-CoA. Altogether, this might be indicative of an overall lower glycolytic activity in the *KCNJ16*^-/-^ kidney organoids.

Lipid droplet aggregation in the kidney and consequent sustained lypotoxicity have been correlated with kidney fibrosis and eventually chronic kidney disease [39, 46–48]. Despite kidney fibrosis has yet not been described as a phenotype for *KCNJ16-* related kidney disease, we observed an increase in the fibrotic markers collagen-I and fibronectin in our *KCNJ16*-depleted kidney organoids when compared to the *KCNJ16*^WT^ control. We hypothesize that kidney fibrosis might appear because of our model itself as our *in vitro* system does not allow for rescue mechanisms to detect or halt the potential damage derived from Kir5.1 loss, and the culture conditions have not been optimized for the need of these cells. Numerous studies have shown that restoring either the defective fatty acid degradation or fatty acid synthesis in kidney cells can mitigate kidney fibrosis progression [49–52]. Statins have gained popularity in pharmacological research because of their high efficacy to reduce lipid accumulation and preventing disease progression originated from lipid disorders, such as diabetic nephropathy and chronic kidney disease [53–55]. Our results showed that 6-days treatment with non-toxic doses of fatty acid synthase inhibitors, especially the combination of simvastatin and C75, was able to inhibit lipid droplet accumulation and subsequently reduce collagen-I deposition. Although exhibiting normal urinary concentrating ability, these patients manifest diverse health complications, including metabolic alkalosis/acidosis, hypokalemia, salt wasting, and sensorineural deafness, significantly influencing their quality of life [2]. Consequently, the potential amelioration or suppression of these symptoms through statins treatment holds considerable importance for both the affected individuals harboring Kir5.1 defects and the broader domain of other lipid-related kidney conditions.

Altogether, these findings confirm that the loss of both healthy alleles of the *KCNJ16* gene negatively impacts and changes the metabolic landscape of kidney organoids towards TCA cycle alterations and induces lipid droplet accumulation. While only biallelic mutations in the *KCNJ16* gene appeared disease causing, we included the *KCNJ16^+/-^* clone to investigate whether monoallelic mutations in *KCNJ16* may express a tubulopathy phenotype as well, which could provide insight into whether carriers of *KCNJ16* mutations are at risk of developing kidney disease. The *KCNJ16*^+/-^ organoids showed a high variability response to changes in pH and overall metabolism were similar to *KCNJ16*^WT^, confirming that only biallelic mutations cause tubulopathy while having one healthy allele remaining seems to show an ameliorated version, still comparable to the wild-type phenotype.

## Conclusions and future directions

In this study, we generated a novel kidney organoid model using CRISPR/Cas9 technology to investigate the recently described *KCNJ16* kidney tubulopathy, confirming the functional impact of *KCNJ16* mutations and revealing metabolic impairment and lipid droplet accumulation in the compound heterozygous mutated organoids. Our findings highlight the importance of kidney organoids and CRISPR/Cas9 technology for the *in vitro* validation of candidate genes, further studying disease phenotypes, and evaluating potential treatment options. While our model provides a valuable tool for understanding the disease phenotype and aid to find therapeutic strategies, future directions should focus on validating these findings in *in vivo* models to further explore lipid accumulation as a novel potential marker for disease detection and to evaluate further therapeutic interventions to improve patient outcomes. Conjointly, these efforts will contribute to advancing our understanding of *KCNJ16* related kidney disease and aid in the development of personalized approaches for diagnosis and treatment.

## Materials and methods

### Antibodies and reagents

All reagents were obtained from Sigma Aldrich (Zwijndrecht, The Netherlands) unless specified otherwise. The primary antibodies used in this study were sheep anti-nephrin (#AF4269-SP, R&D systems, Abingdon, United Kingdom) diluted 1:300, mouse anti-E-Cadherin (#610181, BD Biosciences, Mississauga, ON, Canada) diluted 1:300, rabbit anti-GATA-3 (#5852S, Cell Signaling Technology, Leiden, The Netherlands) diluted 1:300, biotinylated-LTL (#B-1325, Brunschwig Chemie, Amsterdam, The Netherlands) diluted 1:300, rabbit anti-Kir5.1 (#HPA059563, Bio-Connect B.V., Huissen, The Netherlands) diluted 1:300, mouse anti-ZO-1 (#610966, BD Biosciences, Vianen, The Netherlands) diluted 1:200, sheep anti-fibronectin (#AF1918, R&D systems) diluted 1:400, rabbit anti-collagen-I (#ab34719, Abcam, Amsterdam, The Netherlands) diluted 1:300. Primary antibodies were detected using Alexa-647 Donkey anti-sheep (#A-21206) diluted 1:400, Alexa-568, Donkey anti-rabbit (#A10042) diluted 1:400, Alexa-488 Donkey anti-rabbit (#A-21206) diluted 1:400, Alexa-488 Donkey anti-mouse (#A21202) diluted 1:400, Alexa-405 streptavidin conjugate (#S32351) diluted 1:400, all purchased from ThermoFisher Scientific, Breda, The Netherlands. To measure the intracellular pH, the pHrodo™ Red P35372 probe (Thermofisher) was used.

### Cell lines and cell culture

iPSCs were obtained via a material transfer agreement from The Stem Cell Technology and differentiation Center (SCTC) at the Radboudumc, Nijmegen [56]. The iPSCs were cultured in Essential 8 (E8) medium containing E8 supplement and 100 μg/mL Penicillin-Streptomycin (E8 complete medium, all from GIBCO, Life Technologies, Paisley, United Kingdom) in well plates that were coated with 1% Geltrex (ThermoFischer Scientific). Cells were cultured at 37°C in a humified atmosphere in the presence of 5% CO2. When sufficient confluency was reached, cells were washed twice with 1× Dulbecco’s phosphate buffered saline (PBS, ThermoFisher Scientific) and passed into colonies using 0.5 mM EDTA (ThermoFischer Scientific) for 4 minutes at RT. iPSC colonies were collected by addition of E8 complete medium after which colonies were transferred to fresh geltrex coated well plates. To obtain single cells, confluent iPSCs were washed twice with 1× PBS and dissociated into single cells using TrypLE Select Enzyme (ThermoFisher) for 2 minutes at 37 °C. Dissociation was stopped by addition of E8 complete medium, after which cells were pelleted by spinning down at 8000 x g for 5 minutes. The *KCNJ16^+/-^*line harbors a heterozygous frameshift mutation in the exon 5 of the *KCNJ16* gene while the *KCNJ16^-/-^* line harbors a compound heterozygous frameshift mutation in both alleles of the exon 5 of the *KCNJ16* gene.

### Nucleofection of iPSCs

The *KCNJ16* gene was mutated in healthy iPSCs (*KCNJ16^WT^*) using CRISPR/Cas9 combined with nucleofection. Duplex RNA was generated by hybridizing crRNA containing the *KCNJ16* guiding sequence (‘5-‘3 UGAUGCAACUUAAGAUGGAC, IDT, Leuven, Belgium) with fluorescently labeled tracRNA (ATTO 488 and 550, both from IDT) for 5 minutes at 95 °C in a PCR machine (PCR ThermoCycler, Biorad). Next, ribonucleoproteins (RNPs) were generated by addition of Cas9 (Sigma Aldrich) to the gRNA followed by a 20-minute incubation at room temperature. Finally, electroporation Enhancer (IDT) was added, the final reaction mixture consisted of 1.5 μM gRNA, 750 nM Cas9 and 0.05 ng/μL electroporation enhancer. iPSCs were pre-incubated with 10 μM revitacell (GIBCO) 1 hour prior to nucleofection. Approximately 1.0*10^6^ iPSCs were gathered as single cells as described under ‘cell lines and cell culture’ and resuspended in 100 μL NucleofectorTM Solution V (Lonza, Basel, Switzerland) and added to the reaction mixture, after which the mixture was transferred to an electroporation cuvette and cells were electroporated in a 2b-Nucleofector (Lonza). Finally, cells were seeded in geltrex coated well-plates containing E8 complete medium supplemented with 10 μM revitacell.

### Cell sorting with Fluorescence-activated Cell Sorting (FACS)

Cells were dissociated into single cells 16-24 hours after nucleofection, as described in under ‘cell lines and cell culture’. Collected cells were resuspended in E8 complete medium and transferred through a 40 μm cell strainer (Corning Incorporated, NY, USA) to obtain single cells. Next, cells were sorted into single cells at the Flow Cytometry and Cell Sorting Facility at the Faculty of Veterinary Medicine (Utrecht, the Netherlands). Cells that showed fluorescence in the 488 or 550 (depending on the used tracRNA) spectra were selected and seeded as single cell per well in a Geltrex coated 96-well plate containing E8 completed medium supplemented with Clone-R (STEMCELL Technologies, Vancouver, Canada). Plates were spun down at 400rcf for 1 minute to increase single cell attachment and were expanded at 37 °C in a humified atmosphere in the presence of 5% v/v CO_2_ until colonies were formed that could be used to generate a culturable cell line.

### DNA extraction and PCR

iPSCs were gathered as single cells as described under ‘cell lines and cell culture’. Cell pellets were washed once with 1×PBS after which DNA was isolated using the QIAamp DNA Mini Kit (Qiagen, Venlo, The Netherlands) according to manufacturer’s protocol. Next, the *KCNJ16* gene was amplified with 1× Q5 Hot Start High-Fidelity master mix (New England Biolabs, Ipswich, United Kingdom), 0.5 μM forward primer (‘5-‘3 CTACCCGCCAGAGCACATTAT, ThermoFischer) 0.5 μM reverse primer (‘5-‘3 TCTCGAACTGGTGGTGCTTT, ThermoFischer) and 100 ng template DNA. The resulting PCR product was purified using the QIAquick PCR Purification Kit (Qiagen) according to manufacturer’s protocol. Finally, the PCR products were sent for sanger sequencing to Macrogen Europe (Amsterdam, The Netherlands). CRISPR/Cas9-induced mutations were detected using TIDE web tool (https://tide.nki.nl/).

### Differentiation of iPSCs into kidney organoids

iPSCs were gathered as described under ‘cell lines and cell culture’. and were seeded at a density of (500,000 cells/well) in 1% w/v Geltrex coated 6-well plates in E8 containing 10 μM Revitacell for maximum 24 hours. Differentiation into kidney organoids was performed by following the protocol recently described by Jansen, J et al. [57]. Culture medium was changed to Essential 6 (E6, GIBCO) medium supplemented with 6 μM CHIR99021 (R&D systems). Ureteric bud progenitor cells were generated by exposing the cells to the indicated CHIR99021 concentration for 4 days in E6 medium, after which the cells were exposed for 3 days to 200 ng/ml FGF-9 (R&D systems) and 1μg/mL heparin (Sigma Aldrich) in E6 medium. Metanephric mesenchyme cells were generated by exposing the cells to the indicated CHIR99021 concentration in E6 medium for 5 days and exposing for 2 days to FGF-9 and heparin. After differentiation (day 7) into ureteric bud progenitor and metanephric mesenchyme cells, organoids were generated. Cells were washed once using PBS and dissociated into single cells using 0.05% w/v trypsin for 3 minutes at 37°C. Dissociation was stopped by addition of 2 volumes of DMEM-F12 with 10% FCS (GIBCO) and cells were counted. Per organoid, 100,000 Ureteric bud progenitor cells and 200,000 mesenchymal cells were mixed in E6 medium and spun down three times at 300 x g in 1.5 mL Eppendorf tubes. The obtained cell pellets were transferred using sterile wide bored pipets onto a Transwell filter (Corning or CellQart) and exposed for 1 hour to 5 μM CHIR99021 in E6 after which the medium was changed to E6 containing FGF9 and Heparin. Organoids were cultured for 5 days in E6 supplemented with FGF9 and Heparin, after which the organoids were cultured for 13 days in E6 supplemented with 50 ng/mL bone morphogenetic protein-7 (BMP7, R&D systems) and 10 ng/mL human epidermal growth factor (hEGF, Sigma Aldrich). Medium of organoids was refreshed every 2 days, or every 3 days during weekends.

### Bulk RNA sequencing

For RNA-Sequencing analysis, individual organoids were washed 2 times with ice-cold PBS and subsequently dissociated with 350µL of lysis buffer (RLT Buffer, QIAGEN). Samples were stored at - 80°C until they were delivered to USEQ Utrecht Sequencing Facility (Utrecht, the Netherlands). The provided raw transcript counts were further processed with the web application IDEP version 1.12 [58] to generate all the plots shown in the results of this manuscript. The raw data (read counts) was loaded into the IDEP1.12 webtool, normalized by counts per million (CPM) and pre-processed by filtering out genes with a lower than 0.5 CMP. The raw data was then transformed for clustering using the Edge:log2 (CMP + c) method. The heatmap was generated using the hierarchical clustered analysis using Pearson as measure of distance. The differential gene expression (DEG1 and DEG2) was analyzed using the DeSeq2 method with a false discovery rate (FDR) cutoff of 0.5 and a minimum fold-change of 1.5. From the DEG1 and DEG2, the scatterplot, the pathway tree, and the pathways networks were generated. For the pathway network analysis, the absolute values of fold changes were used for the GAGE method, sorting the pathways by both KEGG and molecular function, setting the FDR cutoff at 0.5. Lastly, the KEGG data base and Pathview were used to integrate the up- and down- regulated genes within each available KEGG pathway. The access to the raw data of the RNA sequencing can be obtained upon request.

### Compound testing

All compounds were dissolved in DMSO unless stated otherwise. Since all drugs are already in common use, their dosage for our experiments was determined based on their non-toxic limits in previous literature with similar or same *in vitro* models. For the cyst formation experiments, all organoids were treated at the end of the maturation protocol (d7+18). Organoids were pre-incubated with the corresponding inhibitors for 8h before adding culture media with a fresh solution of inhibitor and forskolin. After 24 hours of treatment, organoids were imaged under brightfield for cyst formation. For counting cysts, RGB brightfield images were converted to 8-bit and circles were drawn around them. The images were blinded, and the cysts were manually counted. For the statins experiments, all organoids were treated 6 days before the end of the differentiation and maturation protocol (d7+12) and media with fresh compounds was replaced daily to overcome the potential fast turnover of the compounds. Forskolin was added at a concentration of 10µM, Digoxin at 500nM, TEA (tetraethylammonium chloride) at 500nM, TGN-20 (2-Nicotinamide-1,3,4-thiadiazole) at 10µM, and Acetamide at 100µM. Simvastatin and pravastatin were added at a concentration of 0.5µM while C75 was added at a final concentration of 40µM, and the same concentrations were kept for the combination treatments. All compounds were purchased from Sigma Aldrich and reconstituted with DMSO to a stock concentration of 1mg/mL for safe and stable storage (−20C). To normalize for the different DMSO content amongst treatment conditions, all organoids were kept at 1% DMSO content, including the untreated controls.

### Immunostainings

All stainings were performed on fixated organoids (4% PFA). After completing the maturation protocol, the organoids were washed 3 times with PBS after which the organoids were fixed with 4% PFA solution (PierceTM 16% formaldehyde (w/v), methanol-free, ThermoFisher) on ice for 20 minutes. After incubation, PFA was removed, and organoids were washed 3 times with PBS. Next, organoids were used for stainings, or organoids were stored in 0.1% v/v formalin at 4°C. For antibody stainings, organoids were blocked in blocking buffer (BB) containing 10% v/v donkey fetal serum (GeneTex) and 0.06% Triton-X (ThermoFisher Scientific) in 1× PBS. Next, primary antibodies in BB were added and incubated overnight while gently shaking at 4 °C. After incubation, primary antibody solutions were removed, and organoids were washed 3 times with 1× PBS for 10 minutes while gently shaking at room temperature. Subsequently, secondary antibodies were added in 0.3% v/v triton- X in 1× PBS and incubated for 3 hours while gently shaking at room temperature. Next, secondary antibody solutions were removed, and organoids were washed 3-4 times with 1× PBS for 10 minutes while gently shaking at room temperature. Finally, organoids were mounted in Prolong gold containing DAPI or without DAPI (Cell Signaling Technology, Leiden, The Netherlands). Images were captured using the confocal microscope Leica TCS SP8 X (Leica Biosystems, Amsterdam, The Netherlands). Senescent probe staining was performed using the CellEventTM Senescence Green Detection Kit (InvitrogenTM) following the manufacturer’s instructions. Lipid droplets were detected using 4,4-Difluoro-1,3,5,7,8- Pentamethyl-4-Bora-3a,4a-Diaza-s-Indacene (BODIPY) probe staining as instructed by the manufacturer. After incubation, organoids were washed 3 times with 1× PBS for 10 minutes while gently shaking at room temperature. Finally, organoids were mounted in Prolong gold containing DAPI. Images were captured using the confocal microscope Leica TCS SP8 X (Leica Biosystems, Amsterdam, The Netherlands).

### Intracellular pH

Mature organoids at d7+18 were dissociated with trypsin, neutralized with E6 media and subsequently centrifuged. Cell pellet was then resuspended in E6 media, and 50,000 cells were seeded in 96-well plates (Corning). Cells were allowed to attach and fill the wells for an average of 2-3 days at 37°C. On the day of the experiment, culture media containing the pHrodo™ Red P35372 probe (Thermofisher) was added to the wells and the plate was introduced into the GloMax plate reader (Promega). The first 7 minutes absorbance was measured at 560nm every minute for baseline pH calibration, after which the media was replaced with the same media as the start of the experiment but with 10mM NaHCO_3_ added. Absorbance was then measured every minute until 20 minutes total experimental time. Absorbance was corrected by total protein amount (in ng) for each sample, extracted with the Pierce BCA protein extraction kit (Thermofisher). The absorbance measurement is indirectly correlated with the pH, the lower the absorbance, the higher the intracellular pH, and vice-versa.

### Swelling assay - Cyst formation

Mature organoids were imaged under brightfield, after which organoids were harvested for stainings as described above. For counting cysts, RGB brightfield images were converted to 8-bit and circles were drawn around them. The images were blinded, and the cysts were manually counted.

### LC-MS based metabolomics

For metabolomics analyses, 48h before sample collection, the normal organoid culture media was substituted by glutamine-deficient media (Gibco, # 11960044) with added 2mM [U-^13^C]-labeled glutamine tracer (Cambridge Isotope Laboratories). We seeded 1 organoid in each transwell insert of a 12-well plate. Organoids were washed 2 times with ice-cold PBS and 500µL of ice-cold lysis buffer (methanol/acetonitrile/dH2O at a 2:2:1 ratio) was added allowing for 1 minute incubation. Organoids were mechanically dissociated by pipetting; the lysate was transferred to a 1.5mL Eppendorf and put in a shaker platform at 4 °C for 20 min. The suspension was then centrifuged at 16.000× g for 20 min at 4 °C and the supernatants containing the metabolite suspension were collected and stored at −80 °C until LC-MS measurement was performed. All samples were sent to the Metabolism Expertise Center (Utrecht University, Utrecht, The Netherlands) for LC-MS measurement and analysis. In short, LC-MS analysis was performed on an Q-Exactive HF mass spectrometer (Thermo Scientific) coupled to a Vanquish autosampler and pump (Thermo Scientific). Metabolites were separated using a Sequant ZIC-pHILIC column (2.1 cm × 150 mm, 5 μm, guard column 2.1 cm × 20 mm, 5 μm; Merck) with elution buffers A (acetonitrile) and eluent B (20 mM (NH_4_)_2_CO_3_, 0.1% NH_4_OH in ULC/MS-grade water (Biosolve, Valkenswaard, The Netherlands)). Gradient ran from 20% eluent B to 60% eluent B in 20 min, followed by a wash step at 80% and equilibration at 20%. Flow rate was set at 150 μL/min. Analysis was performed using the TraceFinder software (ThermoFisher Scientific, Waltham, MA, USA). Metabolites were identified and quantified based on exact mass within 5 ppm and further validated by concordance with retention times of standards. Peak intensities were normalized based on total ion count. Data was further analyzed using the publicly available code from MetaboAnalyst 5.0 [59]. The raw data can be accessed in the external links of this manuscript.

### Data analysis

Every experiment was performed in at least three biological replicates, including at least 3 technical replicates each, unless specified otherwise. Results are shown as the mean ± standard error of the mean (SEM). All gathered numeric data was analyzed using GraphPad 9.0 (GraphPad software, La Jolla, CA, USA). For statistical analysis, one-way ANOVA was performed followed by Tuckey post-hoc analysis unless specified otherwise. Image analysis was performed with ImageJ (ImageJ software version 2.9.0/1.53t, National Institutes of Health, Bethesda, MD, USA). For semi-quantitative analysis of fluorescent images, RGB images were converted to 8-bit. Subsequently, grey scale thresholds were adjusted to only represent true staining. Pixel area limited to the threshold was measured for stainings or for DAPI. Eventually, pixel area of stainings was divided by the pixel area of DAPI for each image.

## Author Contributions

Conceptualization, ESG, AMG, RM and MJJ; methodology, ESG, G.S, AMG; software, ESG and GS; validation, ESG; formal analysis ESG; investigation, ESG and GS; resources, MJJ and RM; data curation, ESG; writing—original draft preparation, ESG; writing—review and editing, ESG, AMG, EAZ, JHFB, JGH, RM and MJJ; visualization, ESG; supervision, AMG, RM and MJJ; project administration, MJJ and RM; funding acquisition, JHJ, MJJ and RM. All authors have read and agreed to the published version of the manuscript.

## Acknowledgements

The authors wish to thank the Radboudumc Stem Cell Technology and differentiation Center (https://www.radboudumc.nl/en/research/radboud-technology-centers/stem-cells) for reprogramming and characterizing the control cell line. The authors would also like to thank Sam Biermans for the cell culture support and the valuable discussions.

## Supplementary Figures

**Figure S1.**
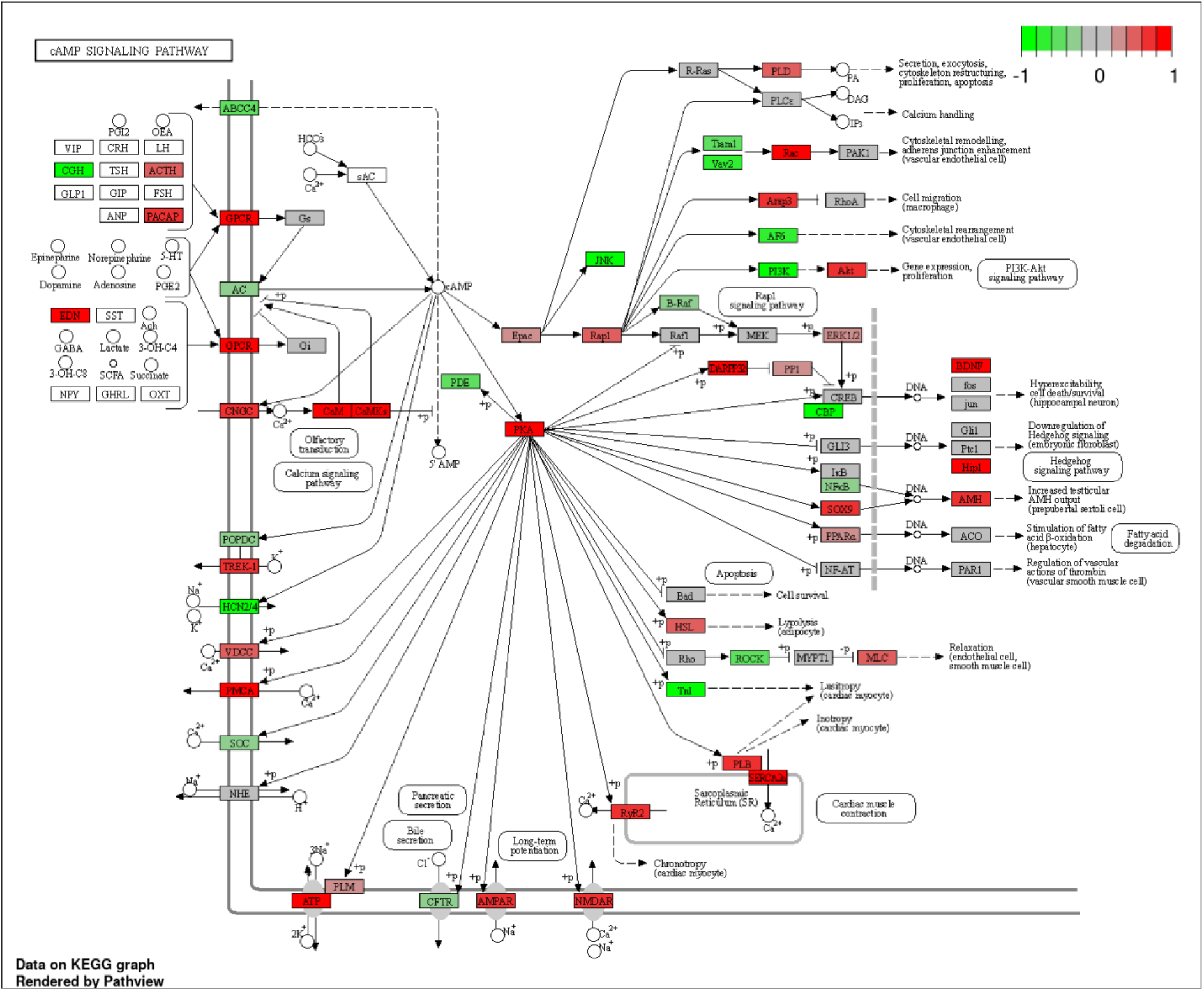
Overview of differentially expressed genes involved in the cAMP signaling pathway. Upon loss of Kir5.1, *KCNJ16^-/-^* kidney organoids showed differences in expression pattern in key genes involved in the cAMP-PKA pathway, when compared to *KCNJ16^WT^*. Upregulated and downregulated transcripts are represented in red and green, respectively. Unchanged transcripts are shown in gray. Transcripts that either could not be measured or did not pass the cutoff thresholds for analysis are shown in white. The upregulated and downregulated transcripts are based on their FDR score.

**Figure S2.**
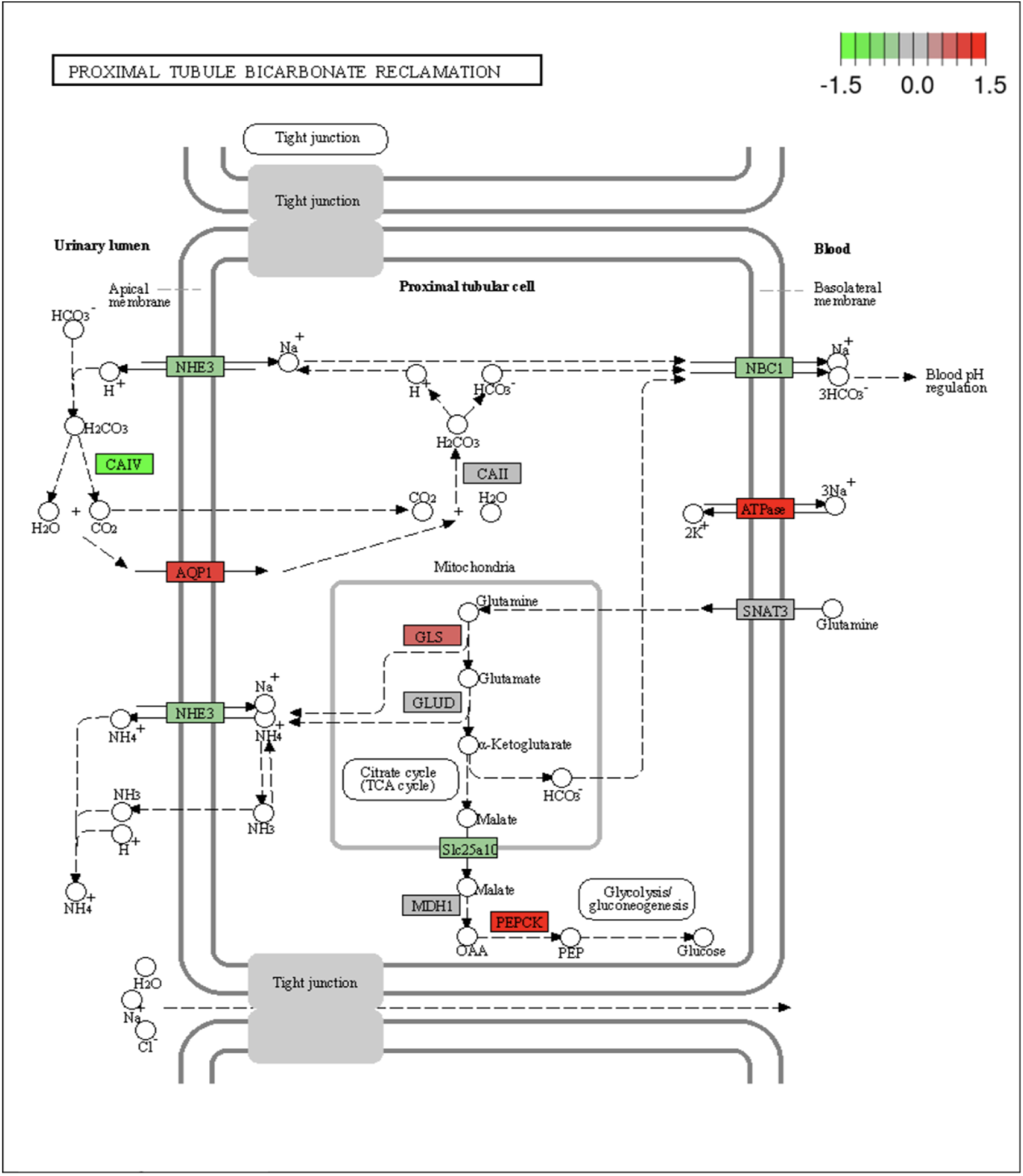
Overview of up- and down- regulated genes involved in the PT bicarbonate reclamation. Upon loss of Kir5.1, *KCNJ16^-/-^* kidney organoids showed differences in expression pattern in key genes involved in bicarbonate reclamation, when compared to *KCNJ16^WT^*. Upregulated and downregulated transcripts are represented in red and green, respectively. Unchanged transcripts are shown in gray. Transcripts that either could not be measured or did not pass the cutoff thresholds for analysis are shown in white. The upregulated and downregulated transcripts are based on their FDR score.

**Figure S3.**
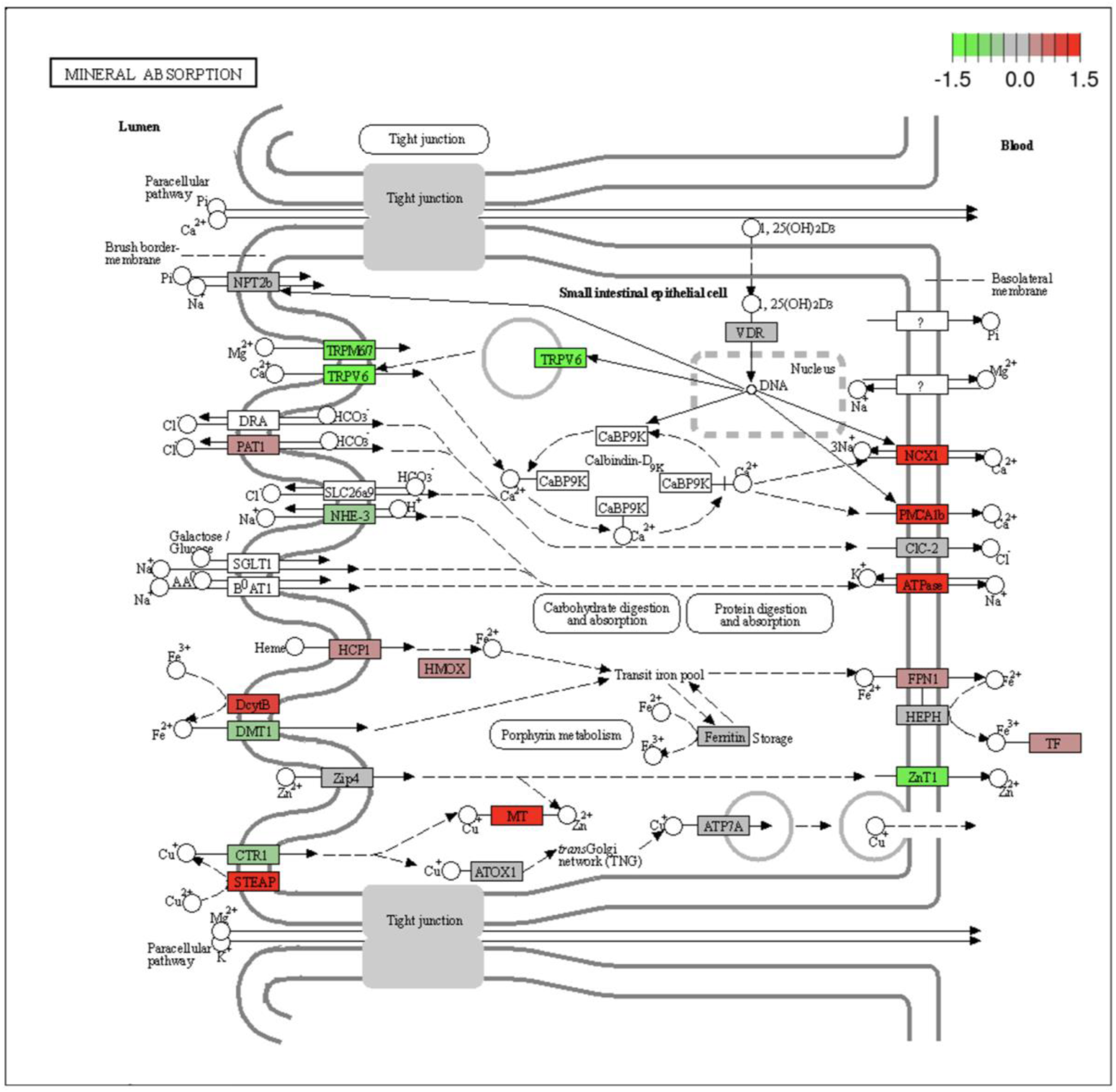
Overview of up- and down- regulated genes involved in mineral reabsorption in the PT. Loss of Kir5.1 resulted in a differential expression pattern in key genes involved in the mineral absorption in the PT. Upregulated and downregulated transcripts are represented in red and green, respectively. Unchanged transcripts are shown in gray. Transcripts that either could not be measured or did not pass the cutoff thresholds for analysis are shown in white. The upregulated and downregulated transcripts are based on their FDR score.

**Figure S4.**
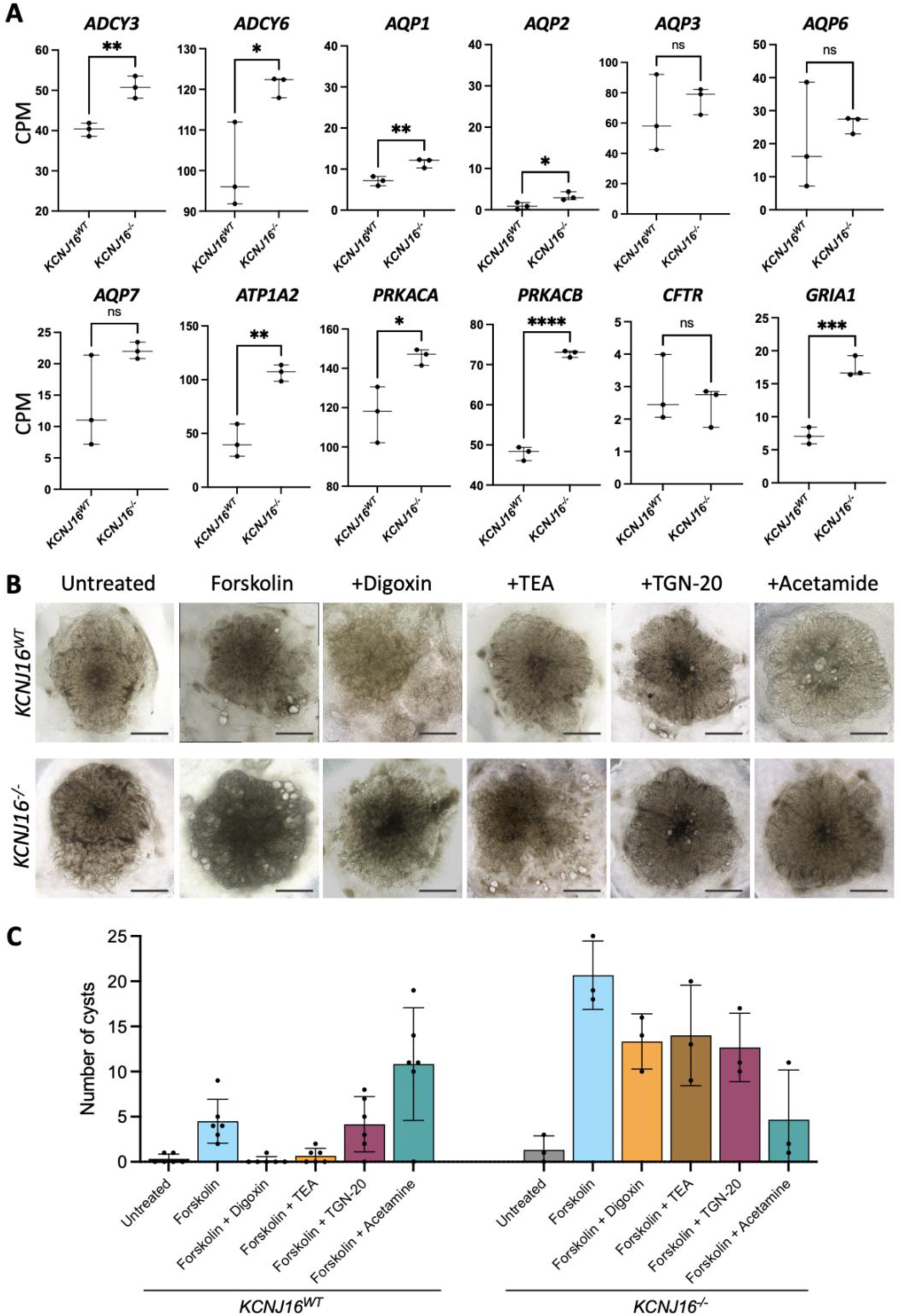
Loss of Kir5.1 results in electrolyte and ion transport impairment. (**A)** The mRNA expression levels quantified as count per million (CPM) of key transporters within the cAMP-PKA pathway. (**B)** Brightfield images of the kidney organoids under untreated and treated conditions with inhibitors of transporters within the cAMP-PKA pathway. (**C)** Graphical representation of the cyst quantification data under untreated and treated conditions with inhibitors of key transporters within the cAMP-PKA pathway

**Figure S5.**
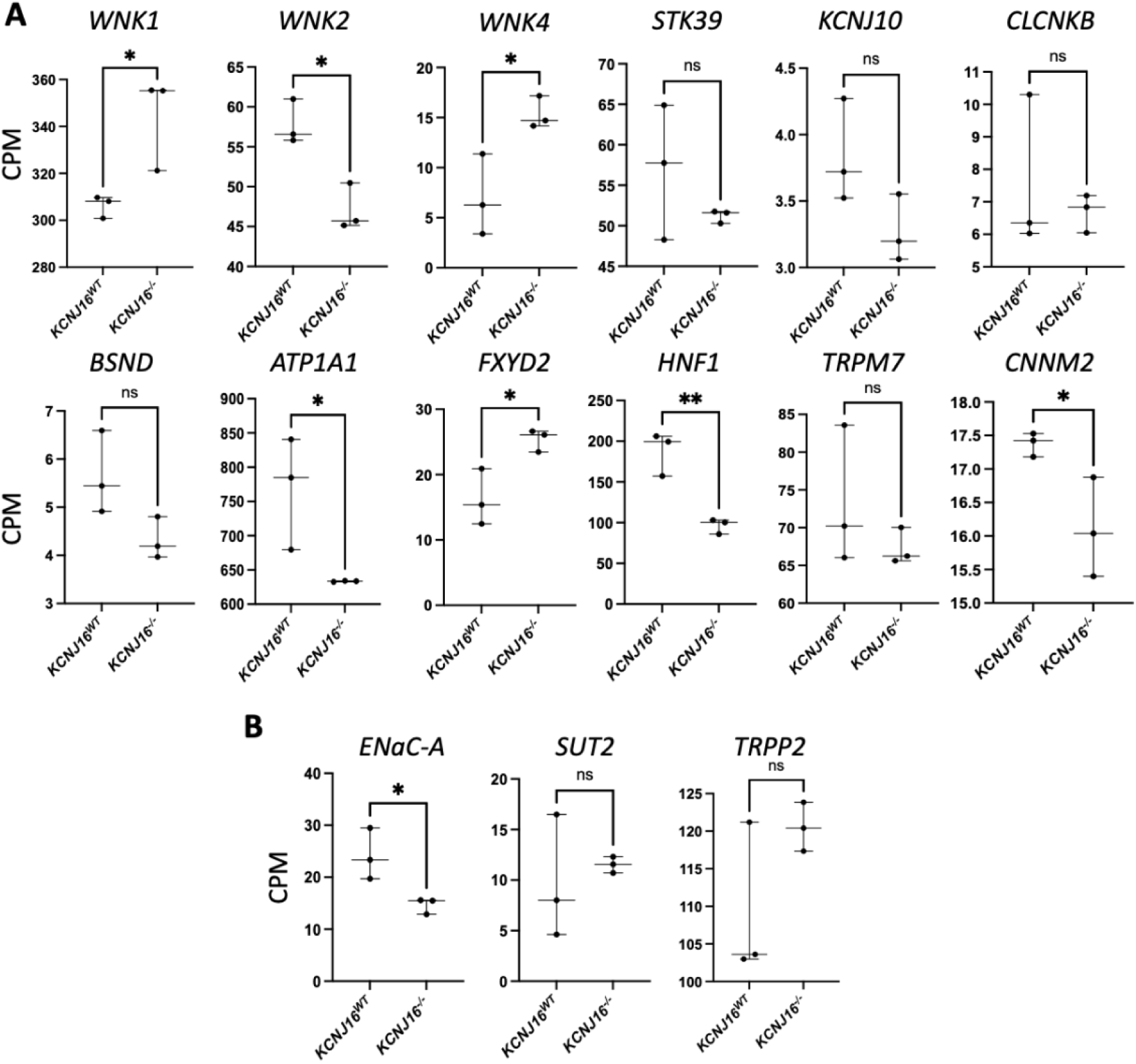
The mRNA levels (CMP) of key genes related to ion and salt transport in the distal part of the nephron. (**A)** The mRNA expression levels quantified as count per million (CPM) of key transporters involved in sodium, potassium, and magnesium regulation in the distal tubule. (**B)** The mRNA expression levels quantified as count per million (CPM) of key transporters involved in compensatory mechanisms upon ion/electrolyte imbalance in the collect duct.

**Figure S6.**
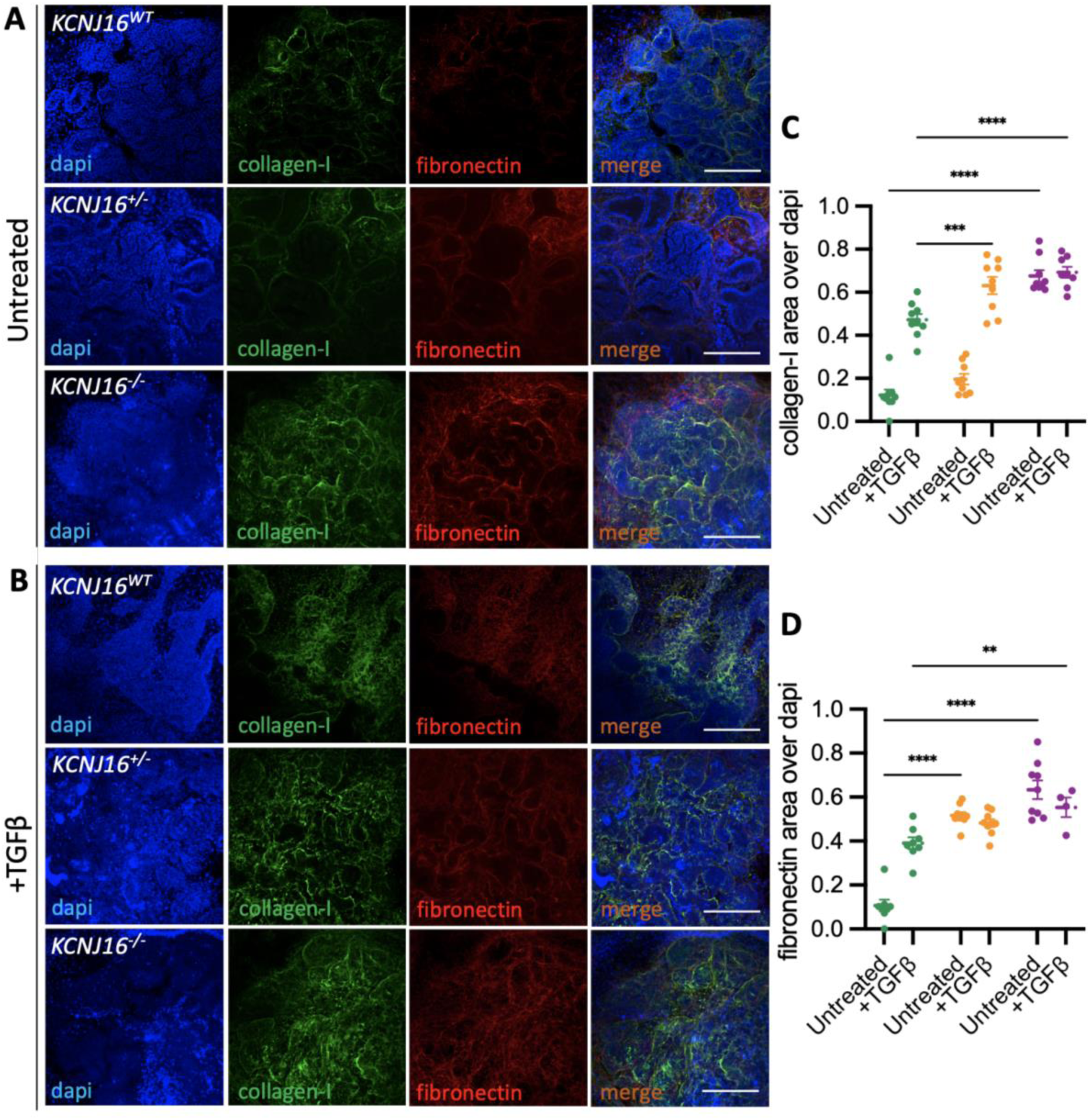
Loss of Kir5.1 promotes fibrosis in kidney organoids. (**A-B)** Immunofluorescence images showing the presence of the fibrotic markers collagen-I and fibronectin in all three *KCNJ16* lines with and without 48h stimulation with TGFß. (**C-D)** Graphical representation of the image analysis shows the accumulation of collagen-I and fibronectin in *KCNJ16*^-/-^ under both TGFß untreated and treated conditions, when compared to the *KCNJ16*^WT^. Scale bars represent 100µm in panels A and B. Statistics represent the significance of one-way ANOVA test (N=3, ****= p<0.0001, ***= p<0.001, **= p<0.01).

**Figure S7.**
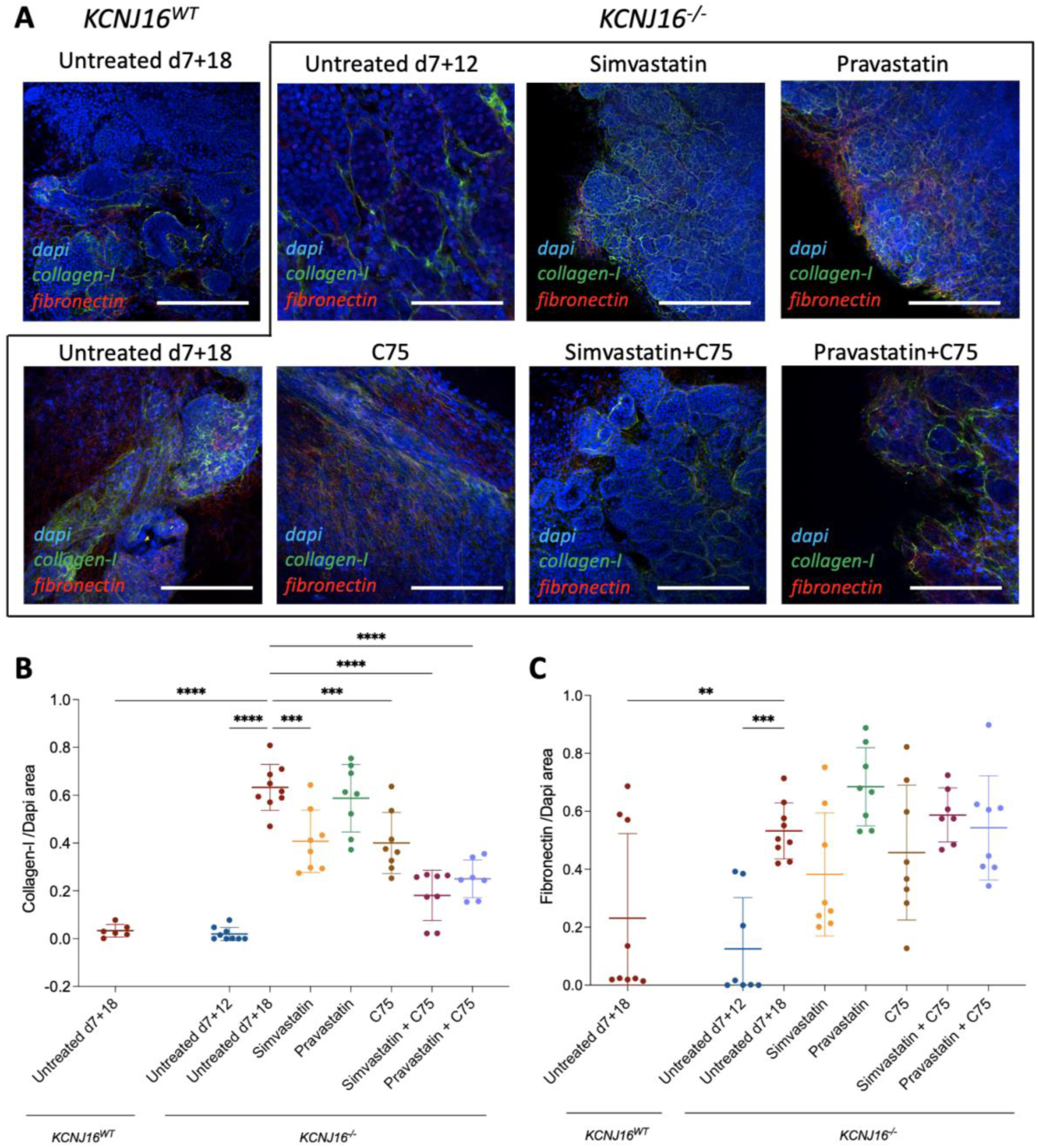
Treatment with statins prevents fibrotic fibers deposition in *KCNJ16*^-/-^ kidney organoids. (**A)** Immunofluorescence images of the fibrosis markers collagen-I and fibronectin in *KCNJ16*^-/-^ organoids after 7 days treatment with simvastatin, pravastatin, C75, and combinations. (**B-C)** Graphical representation of the collagen-I and fibronectin deposition upon treatment with statins. Simvastatin and pravastatin were used at a final concentration of 0.5µM and C75 was used at a working concentration of 40µM. Scale bar 100µm. Statistics represent the significance of one-way ANOVA test (N=3, **= p<0.01, ***= p<0.001, ****= p<0.0001 when comparing all conditions to the untreated *KCNJ16*^-/-^ at d7+18).

